# Intensive and Specific Feedback Self-control of the Argonautes and MicroRNA Targeting Activity

**DOI:** 10.1101/406926

**Authors:** Degeng Wang, Audrey Gill, Fangyuan Zhang

**Author notes:** Current address: Phil and Penny Knight Campus Internship Program, University of Oregon, Eugene, Oregon 97403 (Audrey Gill).

## Abstract

The miRNA pathway consists of three segments – biogenesis, targeting and downstream regulatory effectors. How the cells control their activities remains incompletely understood. This study explored the intrinsically complex miRNA-mRNA targeting relationships, and suggested differential mechanistic control of the three segments. We first analyzed evolutionarily conserved sites for conserved miRNAs in the human transcriptome. Strikingly, AGO1, AGO2 and AGO3 are all among the top 14 mRNAs with highest numbers of unique conserved miRNA sites, and so is ANKRD52, the phosphatase regulatory subunit of the recently identified AGO phosphorylation cycle (AGOs, CSNK1A1, <u>ANKRD52</u> and PPP6C). The mRNAs for TNRC6, which acts together with loaded AGO to channel miRNA-mediated regulation actions onto specific mRNAs, are also heavily miRNA-targeted. Moreover, mRNAs of the AGO phosphorylation cycle share much more than expected miRNA binding sites. In contrast, upstream miRNA biogenesis mRNAs do not display these characteristics, and neither do the downstream regulatory effector mRNAs. In a word, miRNAs heavily and directly feedback-regulate their targeting machinery mRNAs, but neither upstream biogenesis nor downstream regulatory effector mRNAs. The observation was then confirmed with experimentally determined miRNA-mRNA target relationships. In summary, our exploration of the miRNA-mRNA target relationship uncovers intensive, and specific, feedback auto-regulation of miRNA targeting activity directly by miRNAs themselves, *i.e.*, segment-specific feedback auto-regulation of miRNA pathway. Our results also suggest that the complexity of miRNA-mRNA targeting relationship – a defining feature of miRNA biology – should be a rich source for further functional exploration.

## Introduction

MicroRNAs (miRNA), an evolutionarily conserved category of non-coding RNAs, are vital transcriptome regulators. Components and segments of the miRNA pathway – biogenesis, targeting and regulatory actions – are well-understood. However, how cells control the pathway is not clear yet, and complexity of the miRNA-mRNA target relationship is a challenge.

Most miRNA biogenesis starts from RNA polymerase II production of pri-miRNA transcripts. The Drosha RNase III enzyme, with help from the RNA binding protein DGCR8, processes the pri-miRNA into pre-miRNAs, with one pri-miRNA producing up to 6 pre-miRNAs. Some pre-miRNAs are generated directly from mRNA introns and, thus, bypass the pri-miRNA and Drosha processing steps. The pre-miRNAs move, mainly via the Exportin-5 (XPO5) nucleocytoplasmic shuttle, out of nucleus into cytoplasm. The Dicer1 RNase III enzyme, in collaboration with the RNA binding protein TARBP2, processes the pre-miRNA into a mature 22-nucleotide long miRNA (1-3). The miRNA is then loaded onto the Argonaute (AGO) proteins to exert regulatory actions onto target mRNAs via base pairing between its seed sequence, which in human is only 6-8 nucleotide long, and cognate binding sites.

The loaded AGOs, together with the p-body (processing body) scaffold protein TNRC6 (Trinucleotide Repeat Containing 6) they recruit, form the core of the miRNA targeting machinery, bridging upstream miRNA biogenesis to downstream regulatory effectors. The AGO PAZ domain binds to the 3’-end of loaded miRNA, and the PIWI domain to the 5’-end, orienting the miRNA to facilitate base pairing with target mRNAs. Meanwhile, loaded AGOs disassociate from TARBP2 and DICER1 (3), and recruit TNRC6A/B/C. The TNRC6s, in turn, recruit downstream effectors – general translation inhibition and/or mRNA destabilization machinery such as the CCR4-NOT and PAN2-PAN3 complexes. Thus, the AGOs, together with recruited TNRC6s, channel miRNA-mediated regulatory actions onto specific target mRNAs. Given such importance, it is not surprising that AGOs are regulated by many post-translational mechanisms (4). For instance, there are multiple phosphorylation sites and multiple cognate protein kinases (4-8).

Recently, Golden et al discovered an AGO phosphorylation cycle (9), which was soon independently confirmed in both human cells and in *C. elegans* (10), revealing a new layer of regulation of miRNA targeting activity. Through iterative rounds of CRISPR/Cas9 library screening for regulators of miRNA pathway, they identified ANKRD52 and PPP6C – interacting components of protein phosphatase 6 (PPP6) complex – and cognate protein kinase CSNK1A1. Briefly, miRNA-mRNA binding triggers CSNK1A1 phosphorylation of AGO2 on multiple serine residues (S824 – S834), which are evolutionarily conserved in all AGO proteins and are within a structurally unresolved loop of the PIWI domain near the miRNA-target interface (9,10). The phosphorylation disrupts AGO-miRNA binding to mRNAs (10). Meanwhile, PPP6 de-phosphorylates AGO2, presumably getting it ready for the next target binding and phosphorylation cycle. Thus, the AGOs, ANKRD52, PPP6C and CSNK1A1 form a functional module within the miRNA targeting machinery (9,10).

The shortness of miRNA seed sequence leads to a complexity in the miRNA-mRNA target relationship. It enables individual miRNAs to target multiple, and sometimes a large number of, mRNAs; conversely, one mRNA can have binding sites for potentially a large number of unique miRNAs. Unfortunately, the shortness also means low signal-to-noise ratio in transcriptome-wide miRNA binding site identification efforts. Though somewhat mitigated by analyzing evolutionary conservation or combinatorial patterns of multiple miRNA binding sites (11-13), this technical difficulty is perhaps why the complexity remains largely under-appreciated. Current research focuses mostly on individual binding sites instead of the overall binding site distribution pattern, such as the study of miRNA functions in cell cycle regulation (14,15) and in cancers (16,17).

The miRNA binding site distribution is, however, likely a rich source for functional exploration. In transcription regulation, the notion of functionally related genes sharing common transcription factor (TF) binding sites has long been a fruitful assumption (18,19). It is conceivable to assume the same in miRNA-mediated transcriptome regulation. Moreover, TF binding site distribution often reveals feedback regulation. It can be a TF binding to its sites within its own genomic regulatory regions to feedback-control transcription (20). There are also numerous reports of a miRNA pairing with a TF in a feedback loop; the miRNA regulates the TF mRNA, and the latter regulates the miRNA gene’s transcription (17,21,22). A computational study has shown enrichment of such loops in the human regulatory network (23). As for miRNA pathway itself, DICER1 is directly feedback-controlled by miR-103/107 and Let-7 in human (24,25). And, in *C. elegans*, the alg-1 AGO ortholog is directly regulated by mir-71 (26). However, it is unclear to which extent the pathway is directly feedback-controlled in this manner.

Thus, this study explored the distribution of evolutionarily conserved miRNA binding sites in the human transcriptome and, fortunately, shed new light onto cellular control of the miRNA pathway. Among heavily miRNA-targeted mRNAs, *i.e.*, the mRNAs with highest miRNA binding site counts, we detected significant enrichment of miRNA targeting machinery mRNAs, but neither upstream miRNA biogenesis nor downstream effector mRNAs. The AGO phosphorylation cycle mRNAs best exemplified the enrichment, and also share more than expected common miRNA binding sites. Thus, our analysis suggests intensive and specific auto-feedback regulation of miRNA targeting activity by miRNAs themselves. The results also imply miRNA binding site distribution as a rich resource for further functional exploration.

## Results

### The miRNA binding site distribution pattern

We recently studied miRNA binding site distribution in the human transcriptome (27). The distribution is long known to be un-even; a small number of miRNAs target extra-ordinarily high numbers of mRNAs, and a small number of mRNAs contain extra-ordinarily high numbers of binding sites (28). Using evolutionally conserved miRNA-target relationships in the TargetScan database (11), we showed that the miRNA binding site distribution pattern can be described quantitatively by the so-called scale-free relationship (P_(k)_ ∝ (K+a)^-α^, with P_(K)_ as the number of mRNAs containing K unique binding sites; a and α positive constants) (27), which is commonly seen in many domains of biology such as regulatory networks and protein family size distribution (29-31). This is shown in Figure 1. The TargetScan 7.2 dataset identifies 13,035 mRNAs with at least one conserved 3’-UTR miRNA binding sites. A small number of mRNAs (24, less than 0.2%) are extremely miRNA-targeted, each with 60 or more sites. The vast majority of the mRNAs have much fewer sites; >50% of the mRNAs (6926) have 5 or fewer sites (Fig. 1).

**Figure 1:**
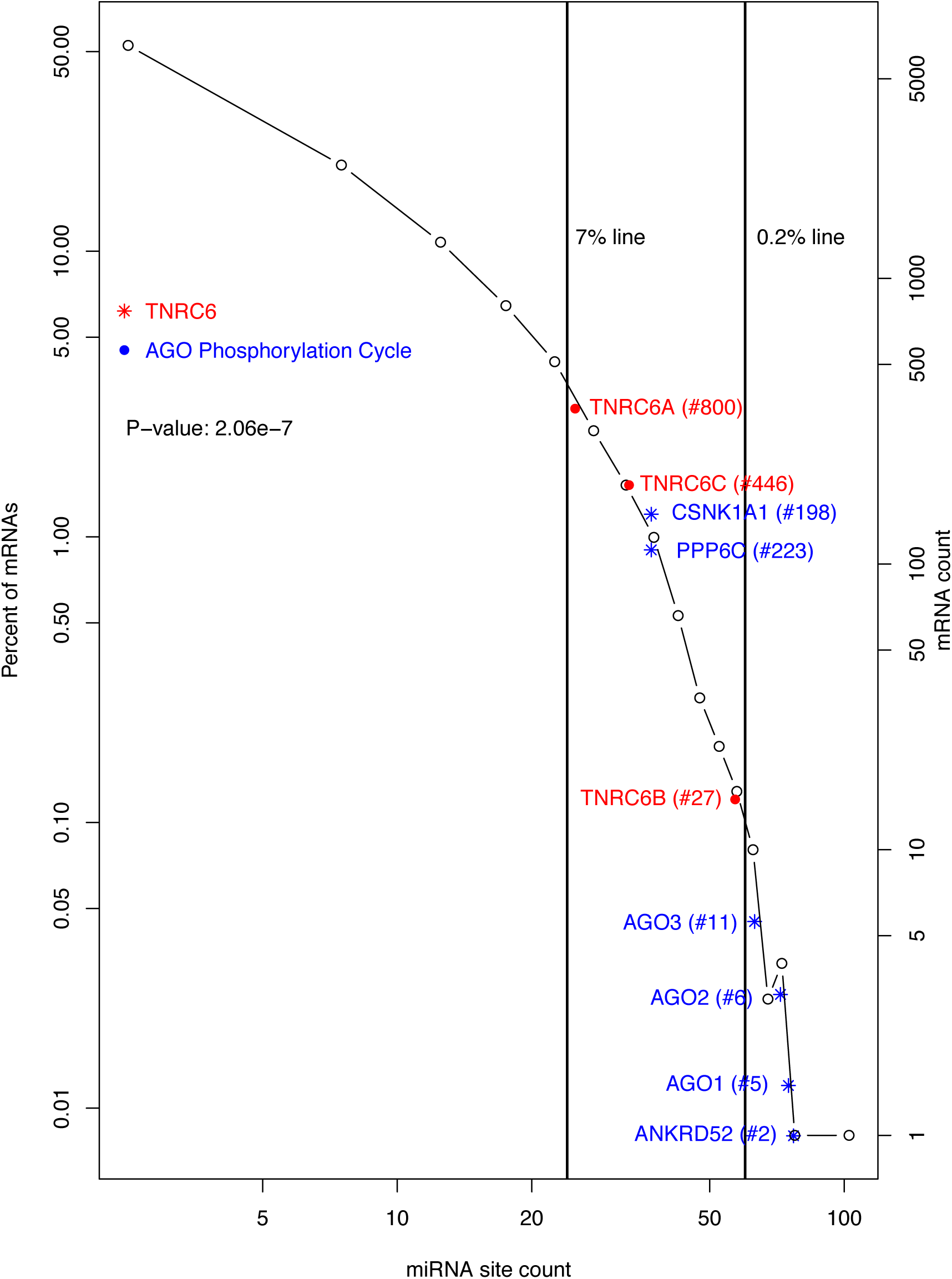
Distribution of miRNA binding sites in the human transcriptome and intensive targeting of AGO phosphorylation cycle and TNRC6 mRNAs by miRNAs. A histogram of human mRNAs based on their miRNA site counts is shown to illustrate the power-law relationship (P_(k)_ ∝ (K+a)^-α^). The two vertical lines denote positions of 0.2 and 7 percentile rankings. AGO phosphorylation cycle (*) and the TNRC6 (•) mRNAs are plotted at their approximate positions to illustrate their high rankings, *i.e.*, intensive targeting by miRNAs. The rankings are given inside the parentheses following gene symbols. The p-value for comparing the whole transcriptome and the miRNA targeting machinery mRNAs (AGO phosphorylation cycle and TNRC6) is specified inside the plot.

### Enrichment of AGO phosphorylation cycle mRNAs among top miRNA-targeted mRNAs

Our purpose in this study was to use the count of conserved miRNA sites as a measurement of the extent to which individual mRNAs are controlled by the miRNAs. We speculated that, if a cellular function was controlled primarily by miRNAs, relevant mRNAs should be enriched among the heavily miRNA-targeted mRNAs, *i.e.*, those with highest conserved miRNA binding site counts. In the case of the miRNA pathway itself, such enrichment implies direct feedback control. Thus, the human mRNAs were ranked based on their binding site counts. We also used the histogram in Figure 1 for schematic interpretation of the site counts of individual mRNAs, *i.e.*, how much a mRNA is miRNA-targeted, relative to the whole human transcriptome.

The top 14 mRNAs with highest miRNA binding site counts in their mRNA 3’-UTRs are listed in table 1, section A. Consistent with previous analyses, TFs and protein kinases are in the list. They also revealed an obvious enrichment of mRNAs for the AGO phosphorylation cycle proteins. Remarkably, AGO1, AGO2 and AGO3 are all in the list. AGO1 is the 5^th^ ranked mRNA, AGO2 the 6^th^ and AGO3 the 11^th^. Even more significantly, ANKRD52 – the mRNA for regulatory subunit of the ANKRD52-PPP6C PPP6 phosphatase complex – is ranked the 2^nd^. This enrichment of AGO1-3 and ANKRD52 mRNAs is illustrated schematically with blue * symbol and text in Figure 1. They are all located to the right of the 0.2% line and, thus, ranked within the top 0.2%, The result strongly suggests that the miRNA regulatory system directly targets the AGO phosphorylation cycle, constituting a major feedback auto-regulatory loop.

**Table 1.**
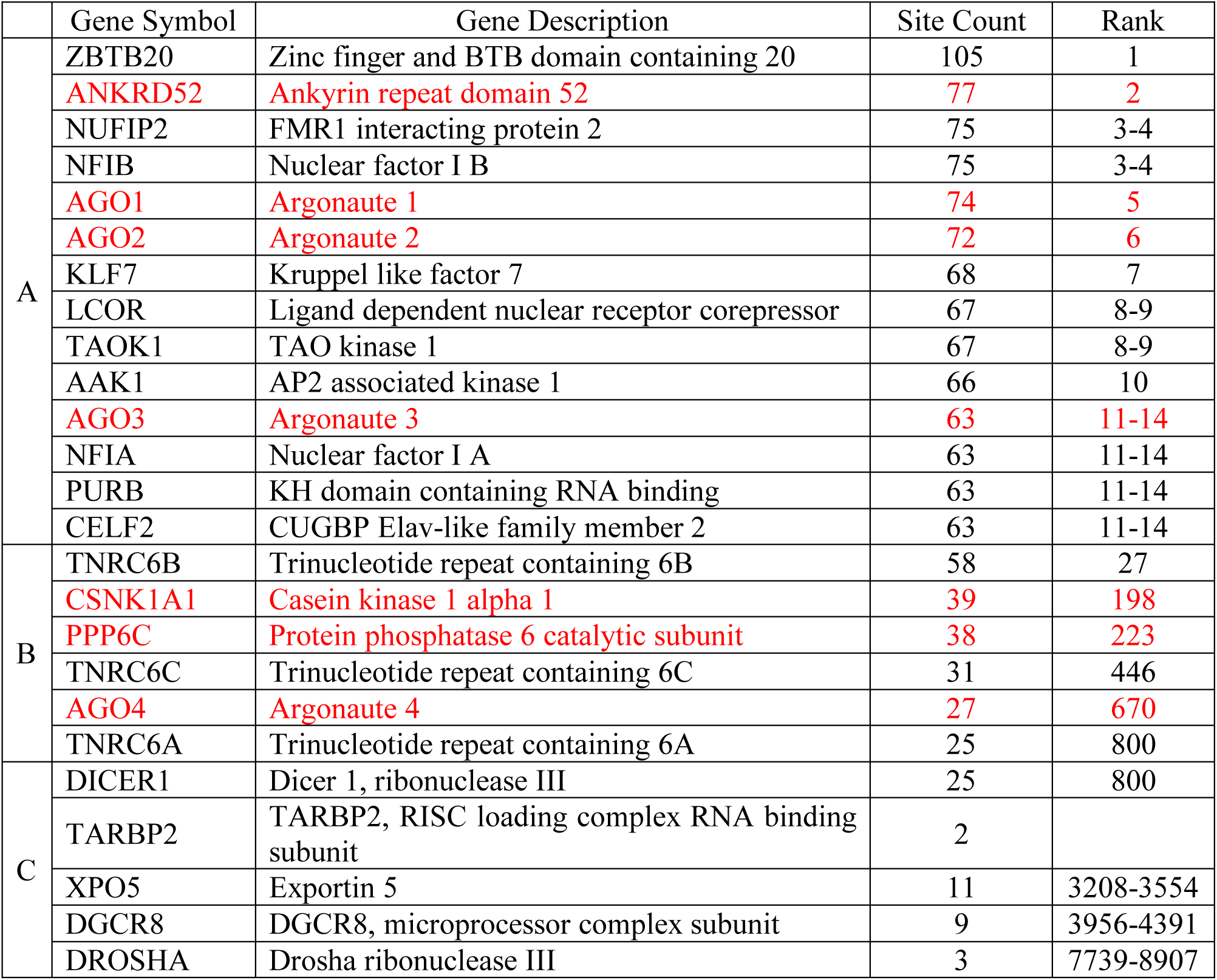
List of the top 14 mRNAs with highest counts of unique conserved 3’-UTR miRNA binding sites (A), other AGO phosphorylation cycle and the TNRC6 mRNAs (B) and miRNA biogenesis mRNAs (C). NCBI gene symbol, description, miRNA site count and descending rank are shown. The AGO phosphorylation cycle mRNAs are highlighted by red font.

|  | Gene Symbol | Gene Description | Site Count | Rank |
| --- | --- | --- | --- | --- |
| A | ZBTB20 | Zinc finger and BTB domain containing 20 | 105 | 1 |
|  | ANKRD52 | Ankyrin repeat domain 52 | 77 | 2 |
|  | NUFIP2 | FMR1 interacting protein 2 | 75 | 3-4 |
|  | NFIB | Nuclear factor I B | 75 | 3-4 |
|  | AGO1 | Argonaute 1 | 74 | 5 |
|  | AGO2 | Argonaute 2 | 72 | 6 |
|  | KLF7 | Kruppel like factor 7 | 68 | 7 |
|  | LCOR | Ligand dependent nuclear receptor corepressor | 67 | 8-9 |
|  | TAOK1 | TAO kinase 1 | 67 | 8-9 |
|  | AAK1 | AP2 associated kinase 1 | 66 | 10 |
|  | AGO3 | Argonaute 3 | 63 | 11-14 |
|  | NFIA | Nuclear factor I A | 63 | 11-14 |
|  | PURB | KH domain containing RNA binding | 63 | 11-14 |
|  | CELF2 | CUGBP Elav-like family member 2 | 63 | 11-14 |
| B | TNRC6B | Trinucleotide repeat containing 6B | 58 | 27 |
|  | CSNK1A1 | Casein kinase 1 alpha 1 | 39 | 198 |
|  | PPP6C | Protein phosphatase 6 catalytic subunit | 38 | 223 |
|  | TNRC6C | Trinucleotide repeat containing 6C | 31 | 446 |
|  | AGO4 | Argonaute 4 | 27 | 670 |
|  | TNRC6A | Trinucleotide repeat containing 6A | 25 | 800 |
| C | DICER1 | Dicer 1, ribonuclease III | 25 | 800 |
|  | TARBP2 | TARBP2, RISC loading complex RNA binding subunit | 2 |  |
|  | XPO5 | Exportin 5 | 11 | 3208-3554 |
|  | DGCR8 | DGCR8, microprocessor complex subunit | 9 | 3956-4391 |
|  | DROSHA | Drosha ribonuclease III | 3 | 7739-8907 |

Next, we examined CSNK1A1 and PPP6C, the respective protein kinase and phosphatase in the AGO phosphorylation cycle. Consistent with our hypothesis, CSNK1A1 and PPP6C are both heavily miRNA-targeted, with 39 and 38 miRNA binding sites, respectively. CSNK1A1 was ranked at the 198^th^, and PPP6C at the 223^rd^, both within the top 2% among the 13,035 genes with at least one conserved 3’-UTR miRNA binding sites. Their high ranks are shown schematically with blue * symbol and text in Figure 1 as well. Thus, six AGO phosphorylation cycle mRNAs (AGO1-3, ANKRD52, CSNK1A1 and PPP6C) are heavily miRNA-targeted, all ranked within the top 2%. It is statistically significant, with a p-value of 1.3e-5 based on a Mann-Whitney-Wilcoxon test. Additionally, though not as heavily targeted as AGO1-3, the AGO4 mRNA has 27 unique 3’-UTR miRNA sites, and ranked at the 670^th^. Thus, the AGO phosphorylation cycle seems under intensive direct feedback regulation by miRNAs.

### The six mRNAs share significant numbers of common miRNA binding sites

It is usually assumed that functionally related genes share similarly gene expression regulation patterns and mechanisms, *e.g.*, regulation by similar sets of transcription factors. In this case, we speculated that the six AGO phosphorylation cycle mRNAs shared more common miRNA binding sites than expected by random chances.

This was indeed the case. The mRNAs for the six proteins share five common miRNA binding sites. This count was significant based on the following analysis. We randomly selected 4 mRNAs from the top 14 mRNAs in table 1, 1 from the 25 genes with 39 miRNA binding sites and 1 from the 22 genes with 30 sites in their 3’-UTRs. We then counted common sites shared by mRNAs of the 6 randomly selected genes. In each computational experiment, this random-selection and calculation process was repeated 1000 times, followed by calculating the proportion of the 1000 attempts that gave a five or higher count. Repeated performance of this computational experiment never generated a proportion higher than 0.05. The proportion fluctuated in a tight range centered at 0.04, suggesting a p-value of 0.04. The significance is also illustrated by a box-plot in Figure 2. The majority of the randomly selected mRNA sets share no or just 1 common miRNA site.

**Figure 2:**
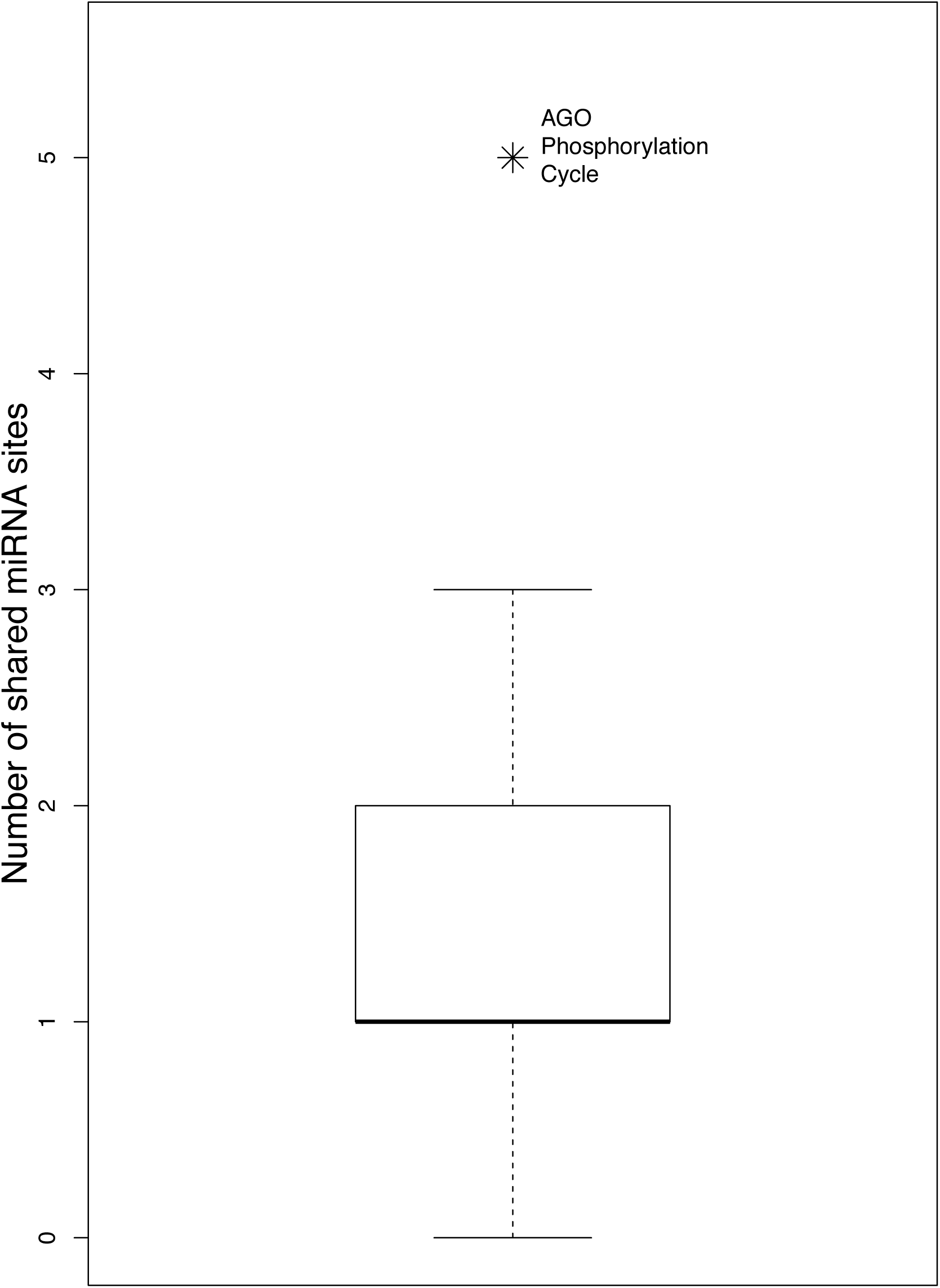
The six AGO phosphorylation cycle mRNAs share more than expected miRNA binding sites. A boxplot of numbers of shared miRNA sites of 1000 sets of randomly selected mRNAs (see text for details) is shown. A symbol is added to illustrate the significance of the observation that the six AGO phosphorylation cycle mRNAs share five sites.

### All miRNA targeting machinery mRNAs are heavily targeted by miRNAs

As discussed in introduction, TNRC6s and AGOs act together to channel miRNA-mediated regulatory actions onto specific mRNAs; AGOs host and position miRNAs for target binding; TNRC6s bridge the loaded AGO proteins to downstream general regulatory effector proteins. Thus, we examined the three TNRC6 mRNAs. Indeed, the TNRC6B mRNA provided clear added evidence for intensive direct feedback regulation of miRNA targeting activity. It has 58 unique miRNA sites, and was ranked the 27^th^ highest among the 13,035 genes. TNRC6A and TNRC6C were also targeted, with 25 and 31 conserved sites, respectively. Their high ranks are also shown schematically with red • symbol and text in Figure 1.

Thus, in terms of the miRNA binding site count, both AGO phosphorylation cycle and TNRC6 mRNAs support intensive auto-feedback regulation of miRNA targeting activity. Statistically, the enrichment has a p-value of 2.06e-7. All of them, including AGO4, are ranked within the top 7% of miRNA-targeted mRNAs, all located to the right of the 7% line (Fig. 1). More significantly, AGO1-3, ANKRD52 and TNRC6B are all among the top 30 mRNAs (Table 1, sections A and B).

### Specificity of the auto-feedback regulation for miRNA targeting activity

We also examined other segments of the miRNA pathway – the upstream biogenesis and the downstream regulatory effectors. No enrichment of miRNA biogenesis mRNAs was detected among the heavily miRNA-targeted mRNAs. The miRNA biogenesis mRNAs (DGCR8, DROSHA, XPO5, TARBP2 and DICER1) are shown in Figure 3A, with red • symbol and text, in contrast to the miRNA targeting machinery mRNAs shown in blue * symbol. None of them had more than 25 miRNA sites, though it has been reported that DICER1 mRNA is directly regulated by miRNAs. Regulation of their expression might be dominated by indirect miRNA feedback control of cognate transcription factor mRNAs as well as other mechanisms (17,21,22,32-34). To some degree, DICER1, the last step of miRNA biogenesis before loading onto the AGO proteins, is a transition point. Its mRNA had 25 unique miRNA sites (ranked at the top 800^th^) – more than its upstream and partner proteins’ mRNAs (DROSHA, DGCR8, XPO5 and TARBP2) but less than miRNA targeting machinery mRNAs (Table 1).

**Figure 3:**
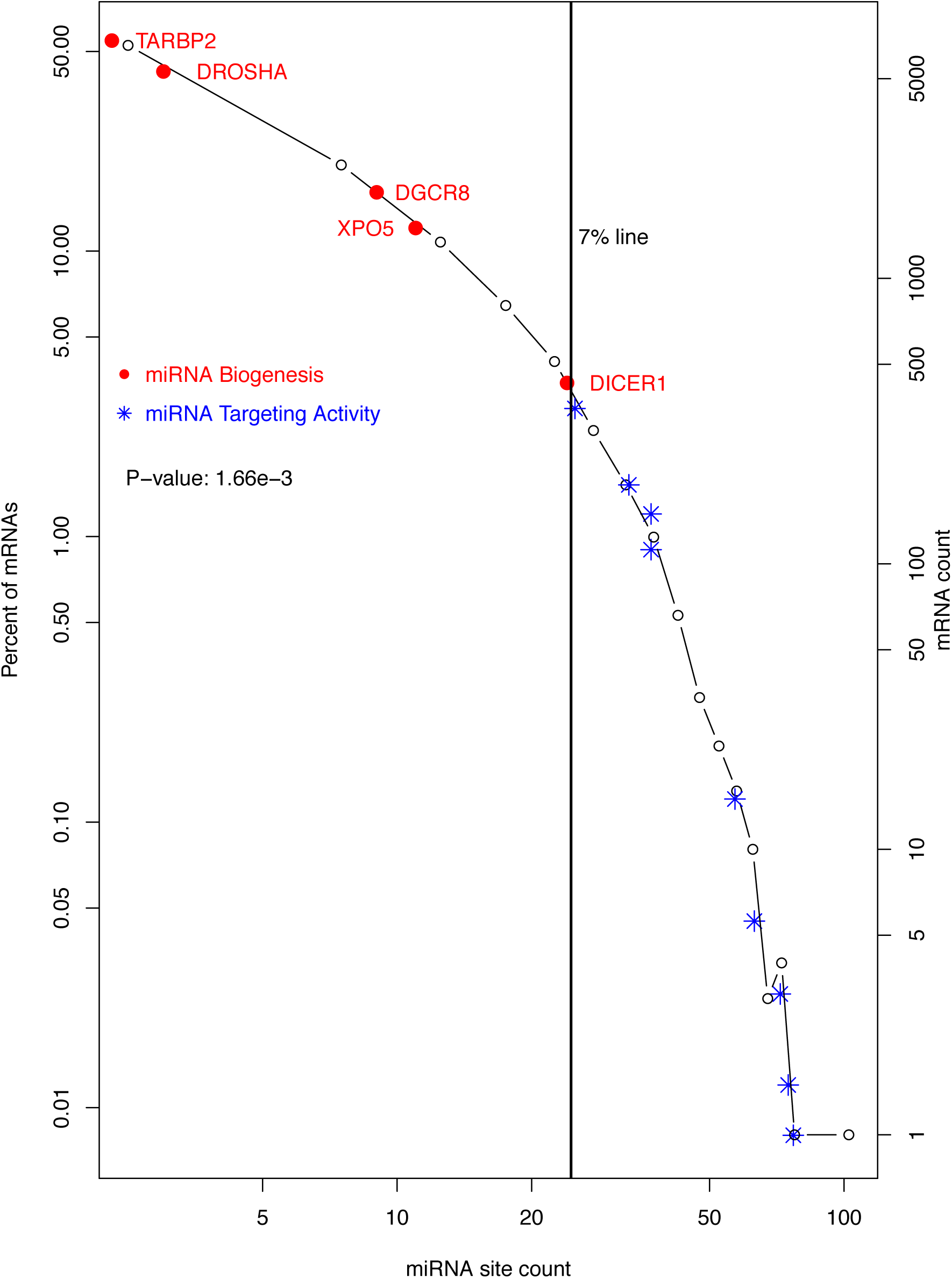
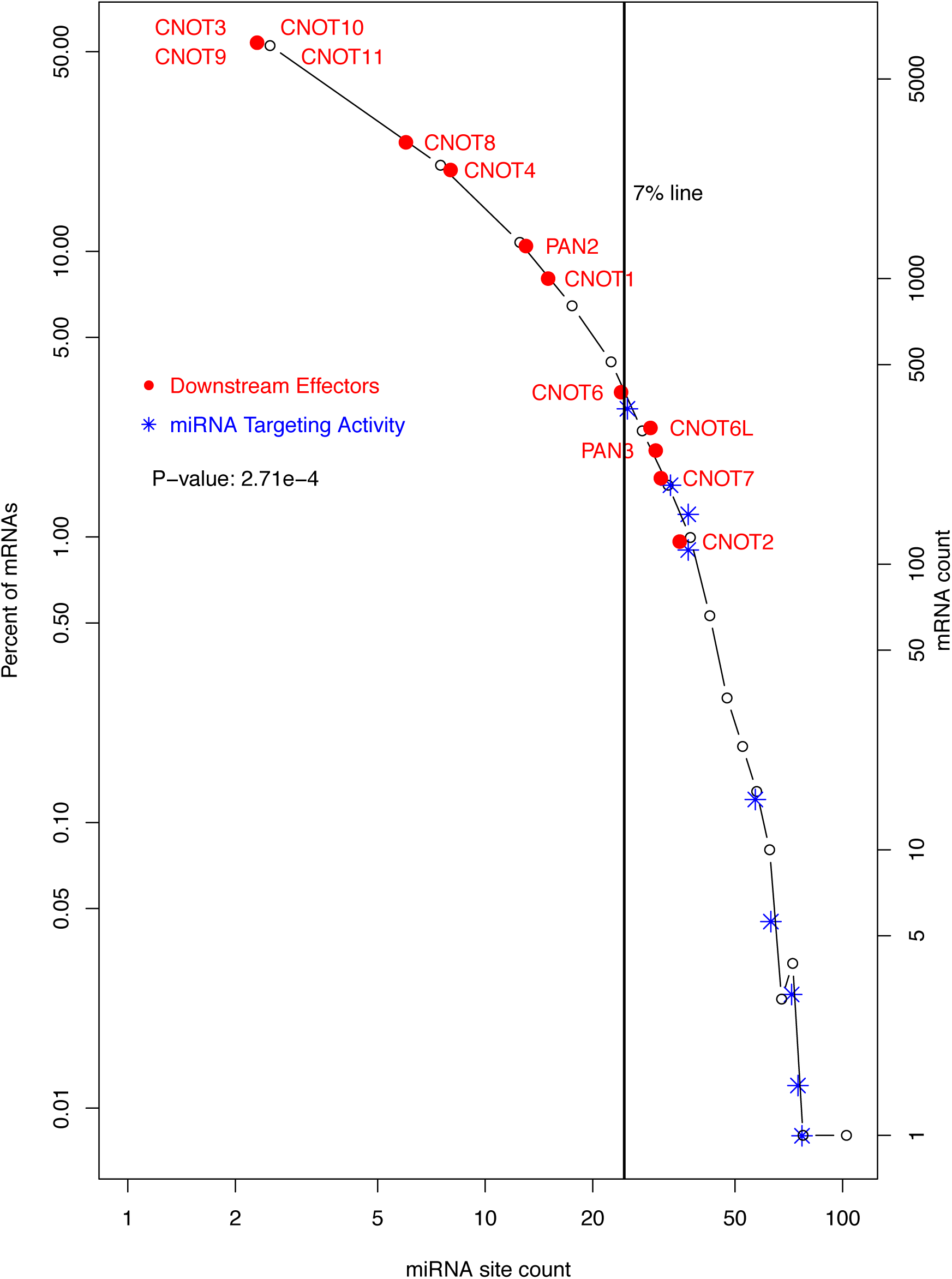
Lower levels of miRNA-targeting of upstream miRNA biogenesis and downstream regulatory effectors. The histogram in Figure 1 is used to show the contrast in the levels of targeting by miRNAs between two groups of mRNAs. The miRNA biogenesis (•) and miRNA targeting machinery (*) are shown in A, and the downstream regulatory effectors (•) and miRNA targeting machinery (*) in B. In B, CNOTs 3 and 9-11 share one data point, as they have the same number of miRNA sites. The p-values of the contrast between two groups of mRNAs are specified inside both plots.

Downstream of the targeting machinery, the general regulatory effector mRNAs also became less miRNA-targeted. This is shown by the CNOTs (CNOT1-4, CNOT6, CNOT6L and CNOT7-11), PAN2 and PAN3 mRNAs schematically in Figure 3B. The CNOT and PAN mRNAs are shown with the red • symbol and text. Compared to the miRNA targeting machinery mRNAs (in blue * symbol), CNOT and PAN mRNAs have much lower miRNA binding site counts.

Thus, the intensive auto-feedback regulation seems specific for the mRNA targeting segment of the miRNA pathway. The specificity is supported by statistical analyses. The miRNA site counts of miRNA biogenesis mRNAs are not significantly different from the whole human transcriptome (p-value 0.33), and neither are those of the effector mRNAs (p-value 0.09). Both the miRNA biogenesis and the effector mRNAs have significantly lower miRNA binding site count then the targeting machinery mRNAs, with p-values of 1.66e-3 and 2.71e-4, respectively (Fig. 3A and B).

The specific auto-feedback regulation of miRNA targeting activity is also illustrated schematically. In Figure 4, the miRNA pathway is divided into 6 steps – DROSHA processing (1), XPO5 nucleus export (2), DICER1 processing (3), the AGO phosphorylation cycle (4), TNRC6 recruitment (5) and downstream regulatory effectors (6). The biogenesis mRNAs (steps 1-3) are shown with black symbols and text; the targeting step mRNAs (steps 4-5) in red symbols and text; and regulatory effector mRNAs (step 6), as exemplified by the CCR4-NOT and PAN2-PAN3 complexes, in blue symbol and text. The black line connects the median miRNA site count for each step, showing that overall miRNA site count increases from the first step of miRNA biogenesis (DROSHA) towards AGO phosphorylation cycle. Subsequently, it decreases towards the TNRC6s, and then towards downstream general effectors. Overall, the counts are higher in the targeting steps (steps 4-5) than in either the biogenesis steps or the downstream effector step (Fig 4).

**Figure 4:**
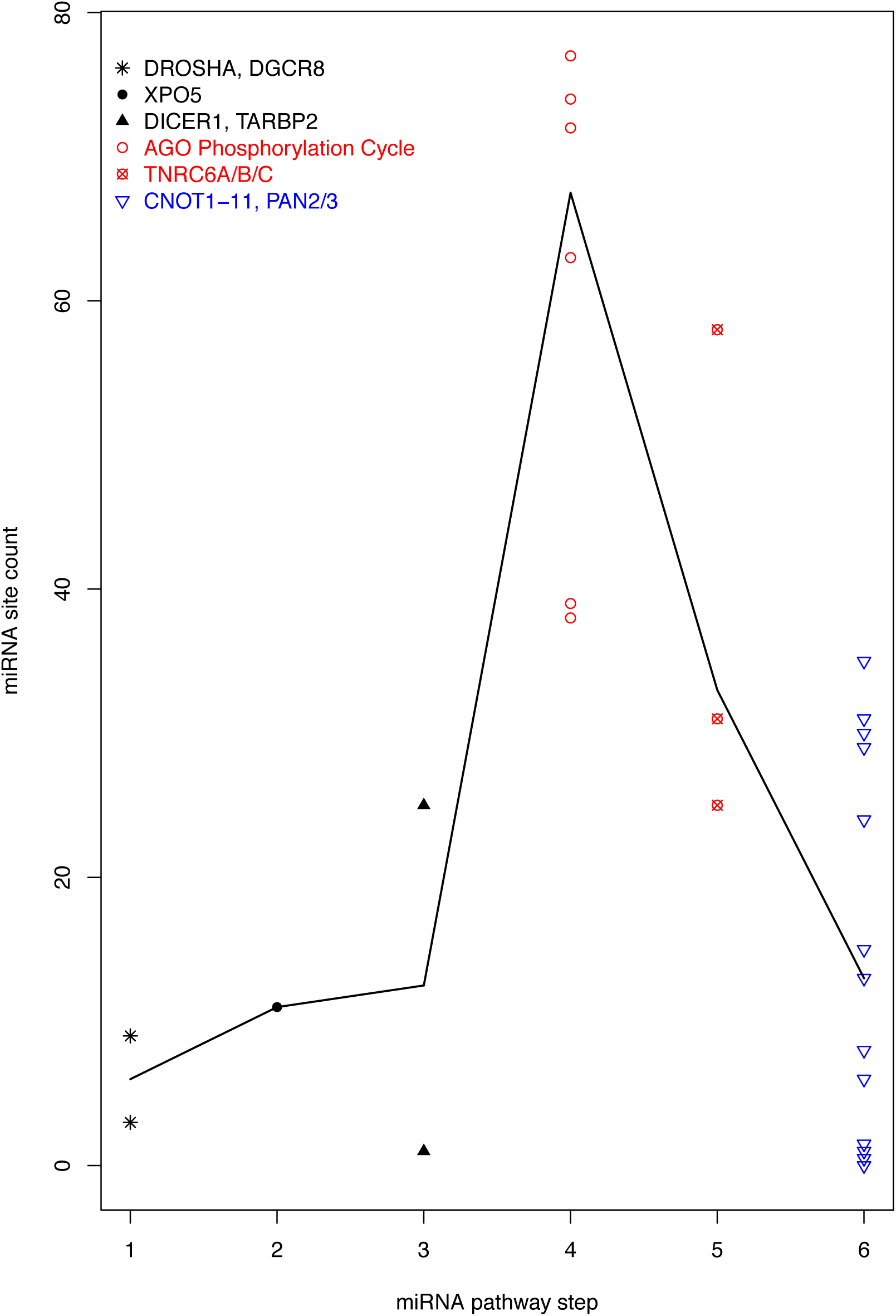
Specificity of auto-feedback regulation for the targeting segment of miRNA pathway. A scatter plot of mRNA miRNA binding site count is shown, with the mRNAs grouped into 6 steps of the miRNA pathway. The biogenesis steps are plotted in black, the targeting steps in red and the downstream effector in blue. The black line connects the median site counts of the 6 steps.

### Confirmation of the enrichment with experimentally determined miRNA binding sites

Another valuable resource for this study is the miRNA-mRNA target relationship identified experimentally with CLIP-seq based high-throughput approaches in a comprehensive and unbiased manner (35-37). The popular miRTarBase database collects and annotates such datasets (38,39). Thus, the miRTarBase CLIP-seq dataset was downloaded. The dataset identified 10,276 mRNAs with at least one miRNA binding sites. We ranked the mRNAs based on their unique miRNA binding site counts in the dataset, and repeated our analysis.

The result confirmed the specific enrichment of miRNA targeting machinery among top miRNA-targeted mRNAs (Fig. 5 and 6). Except for PPP6C mRNA, they all are located to the right of the 15% line and, thus, ranked within the top 15%, (Fig. 5A). AGO2 and AGO3 are both highly ranked, at the 4^th^ and 47^th^, respectively; TNRC6B and TNRC6A at the 40^th^ and 43^rd^, respectively; and so is ANKRD52, ranked within the top 2 percentile. Not surprisingly, the miRNA biogenesis mRNAs are not as highly ranked (Fig. 5A), and neither are the CNOTs, PAN2 and PAN3 mRNAs (Fig. 5B). CNOT3, CNOT8-11 and PAN3 have no miRNA site in miRTarbase dataset (Fig. 5B). Once again, DICER1 was intermediately ranked (Fig. 5A), consistent with its role as a transition point from miRNA biogenesis to target-binding activity.

**Figure 5:**
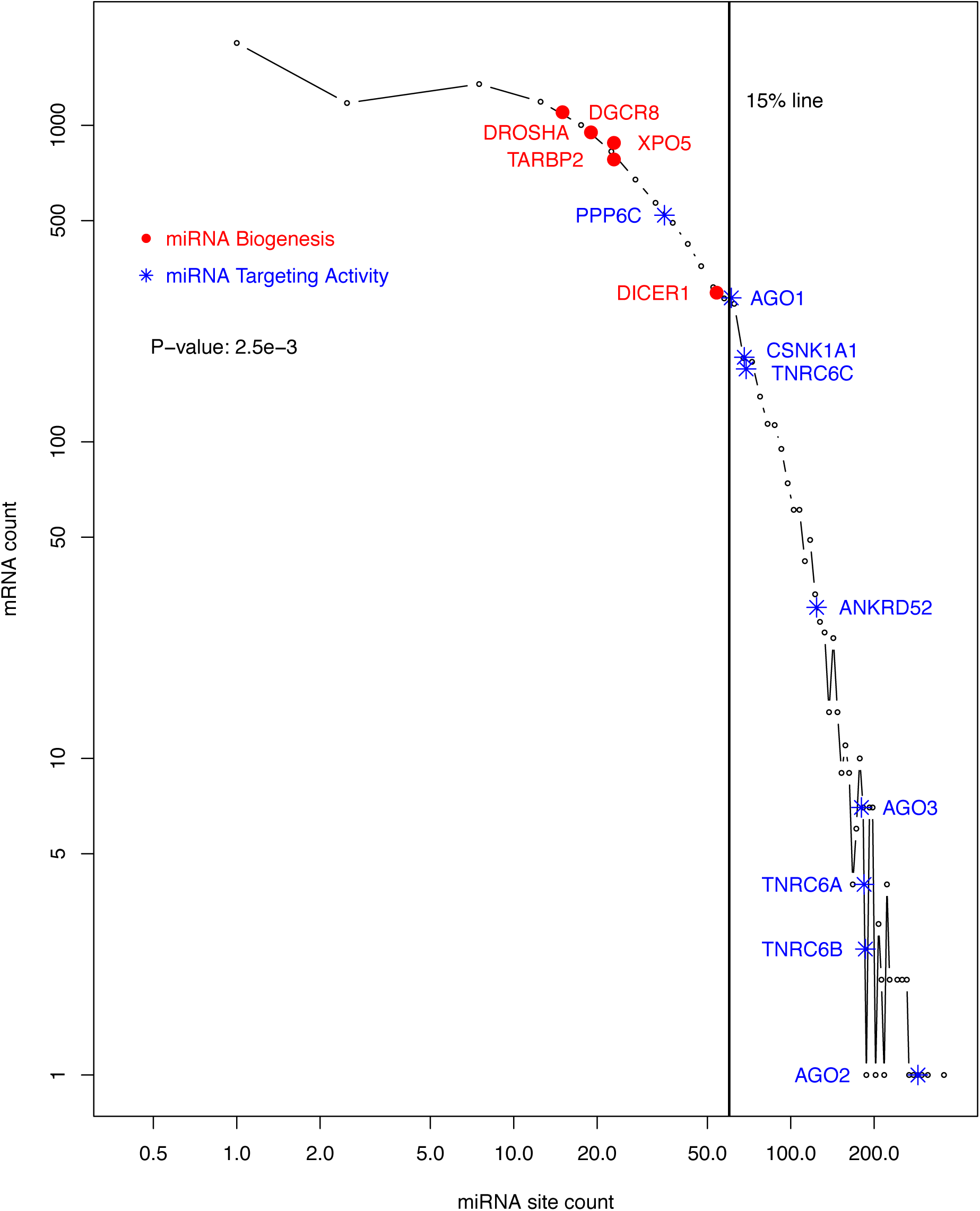
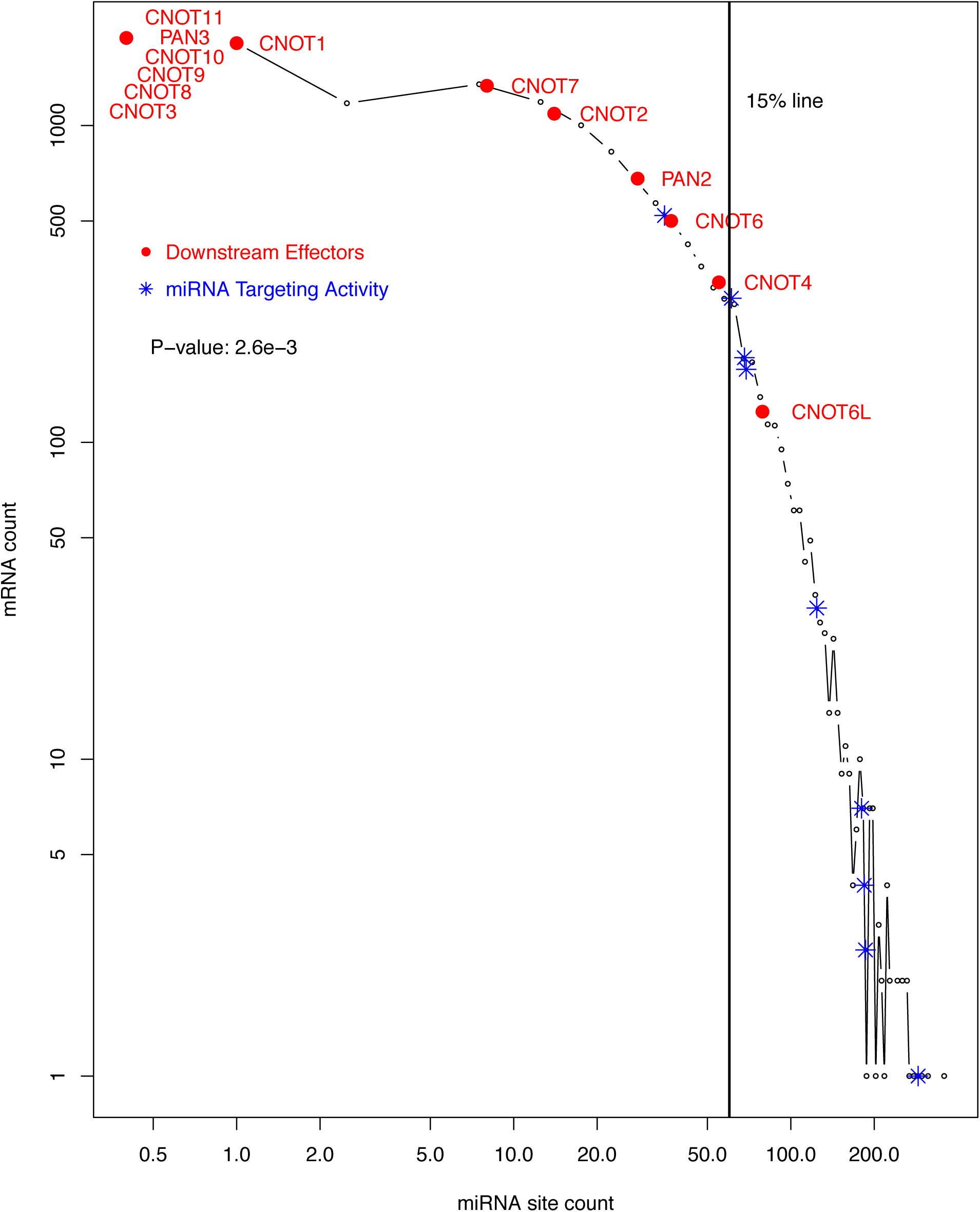
Confirmation of higher levels of miRNA-targeting of the targeting than other segments of the miRNA pathway with the miRTarBase experimental dataset. As in Figure 1, a histogram of human mRNAs based on their miRNA site counts in the miRTarBase dataset is shown, illustrating the power-law relationship. The vertical line denotes the position of 15^th^ percentile ranking. The histogram is also used to display the contrast in the levels of targeting by miRNAs between two groups of mRNAs. The miRNA biogenesis (•) and miRNA targeting machinery (*) are shown in A, and the downsteam regulatory effectors (•) and miRNA targeting machinery (*) in B. CNOTs 3 and 8-11 and PAN3 have no miRNA site in the miRTarbase dataset. They are plotted with one arbitrary data point (B) to help to illustrate the overall lower level of CNOT1-11 and PAN2/3 targeting by miRNAs. The p-value of the contrast between two groups of mRNAs are specified inside both plots.

**Figure 6:**
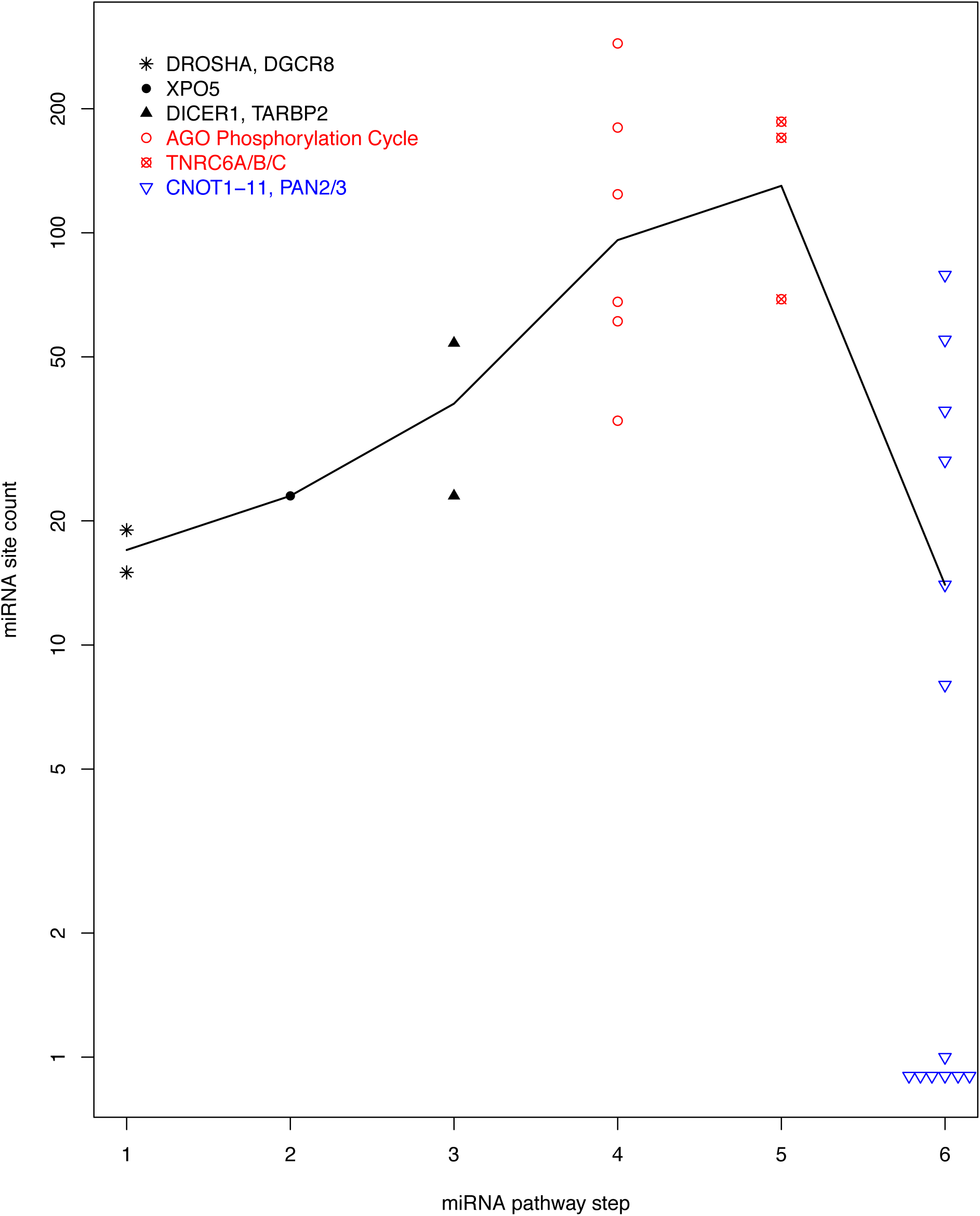
Specificity of auto-feedback regulation for the targeting segment of miRNA pathway based on the miRTarBase experimental dataset. As in Figure 4, a scatter plot of mRNA miRNA binding site count is shown, with the mRNAs grouped into 6 steps of the miRNA pathway. The biogenesis steps are plotted in black, the targeting steps in red and the downstream effectors in blue. The black line connects the median site counts of the 6 steps.

Additionally, the mRNAs are plotted in Figure 6, in the same manner as in Figure 4, by steps of the miRNA pathway. Overall miRNA site count increases from the first step of miRNA biogenesis (DROSHA) towards the targeting steps (AGO phosphorylation cycle and TNRC6s), and then decreases towards downstream general effectors (Fig. 6). The increase of miRNA site counts from miRNA biogenesis to targeting has a p-value of 2.5e-3 (Fig. 5A); the decrease from targeting to downstream effector has a p-value of 2.6e-3 (Fig. 5B).

However, differences were also observed. Overall, the enrichment becames less dramatic; the p-value is very significant, but increases to 0.003 from 2.06e-7. AGO1 and ANKRD52 both ranked high, but not as much as in the ranking based on TargetScan dataset. TNRC6A, on the other hand, became much higher ranked at the 43rd. This might be a reflection of the different biases in the two datasets. The TargetScan dataset is based on computational analysis and, due to technical necessity, covers only orthologously conserved sites. The experimental approaches for the miRTarBase dataset do not have such limitation. Instead, they tend to bias toward miRNAs and mRNAs that are expressed at high levels in the studied cells and under the specific experimental conditions; miRNAs and mRNAs that are not or lowly expressed tend to be missed. To some degree, the biases in the two datasets complement each other. The exact causes for the discrepancy between the two datasets become an interesting topic for our future investigation.

## Discussion

MiRNAs are crucial components of cellular transcriptome regulatory machineries. Tremendous technical challenges remain to be solved. Among them is the low signal-to-noise ratio in miRNA binding site identification due to the shortness of the seed sequences – only 6-8 nucleotides in human. This challenge has been partially alleviated by evolutionary conservation analysis and by detecting combinatorial patterns of multiple sites. Evolutionary conservation analysis is exemplified by the TargetScan dataset. This study took advantage of this dataset and revealed novel insights into how the miRNA pathway itself is regulated. Briefly, the AGO phosphorylation cycle and the TNRC6 mRNAs seems, in terms of their numbers of unique miRNA sites, under intensive direct auto-feedback regulation by miRNAs. This is consistent with frequent auto-feedback regulation of key regulators, such as transcription factors (20). On the other hand, the upstream miRNA biogenesis steps seem not under such regulation. As for the downstream effectors, this regulation seems, though somewhat active, much less dominant. Our observation was confirmed with the miRTarBase experimentally determined dataset as well.

Why the AGO phosphorylation cycle and the TNRC6s are targeted for this direct auto-feedback-regulation is an interesting issue. These proteins link the upstream miRNA biogenesis and the downstream general mRNA inhibition machinery. It is tempting to speculate that this link serve as the rate-limiting step of the miRNA regulatory activity, as such steps generally are under tighter control. However, there seems to be no data available to test this speculation.

Our computational analysis utilized the occurrence of miRNA sites in mRNAs. The sites confer the capacity for the mRNA to be repressed by the miRNA pathway. However, intuitively, this repression should not be constitutive, as the ability to dynamically relieve the repression is operationally advantageous for the cells, *e.g.*, for rapid protein production in case of a need for quick adaptation to signals. Unfortunately, current computational analysis techniques are powerless in discerning such dynamics. To answer this issue, further experimental studies across different physiological conditions and/or transitions and in multiple cell types are needed.

These additional transcriptomic datasets will also enable computational mechanistic investigation into miRNA regulatory actions. While the miRNA sites and other regulatory signals embedded in mRNA sequences are the enabler of the regulatory actions, the datasets describe dynamic mRNA regulation patterns – the results of the regulatory actions. In other word, the datasets will serve as necessary inputs for computational reverse engineering efforts to decode the regulatory signals embedded in mRNA sequences. Among the many questions to be asked is whether and how specific combinatorial patterns of multiple regulatory signals dictate specific mRNA regulation patterns. The pattern can be a combination of multiple miRNA binding sites or a mix of miRNA sites with other regulatory elements, such as mRNA secondary structures.

It should be noted that miRNA-mediated mRNA inhibition exhibit technical similarities with transcription regulation by transcription factors. The complexity of regulator-target relationship applies to transcription regulation as well; a transcription factor usually regulates multiple genes, and a gene is always regulated by multiple transcription factors. And, in both cases, the consensus binding sites are short, leading to low signal-to-noise ratio in computational site predictions (40). Evolutionary conservation helped, in both cases, to alleviate this technical difficulty (41). Combinatorial patterns of multiple transcription factor binding sites and other genomic contextual information have provided additional means for improving the signal-to-noise ratios (42). It seems the combinatorial pattern of multiple sites also apply to miRNA-mediated mRNA inhibition. As our understanding of miRNA regulatory actions improves, it will be interesting to see whether additional similarities exist.

Finally, this study suggests that the overall miRNA binding site distribution pattern should be a rich source for further, technically more sophisticated, functional exploration. This distribution is currently relatively under-appreciated, perhaps due to the high levels of noise in such datasets. However, both computational and experimental approaches will improve, leading to more and more reliable datasets. It is our belief that the binding site distribution will play much more significant roles in our endeavor to a thorough understanding of miRNA-mediated transcriptome regulation.

## Materials and Methods

### Evolutionarily conserved miRNA binding sites

To alleviate the high noise issue associated with computational miRNA binding site prediction, we restricted our analysis to evolutionarily conserved binding sites for conserved miRNA families. The set of sites were downloaded from the TargetScan database 7.2 in July 2018 (11). At the time of download, this was the most current version. The dataset contains 120,702 evolutionarily conserved miRNA binding sites in the 3’-UTRs of 13,035 human genes. We have previously used the dataset of TargetScan 7.1 (27), with which all observations of this study were originally made. During our updating to the new version, we noticed a severe shortening of the AGO2 3’-UTR to 895 base pairs in TargetScan 7.2, which is not consistent with the current AGO2 gene model. So, just for AGO2, we continued to use its information in TargetScan 7.1.

### Experimentally determined miRNA binding sites

We downloaded the miRTarBase release 7.0 (September 2017 release) human data from its websites in July 2018 (38,39). At the time of download, this was the most current version. In this study, only CLIP-seq generated data was used, in order to ensure the data was generated in a comprehensive and unbiased manner. The data was used as a list of experimentally determined miRNA-mRNA target relationship.

### Computer software

The open source software package R (version 3.3) installed on a Mac Pro desktop computer was used for data analysis and plotting. The Mann-Whitney-Wilcoxon tests were performed with the wilcox.test() method.

## Declaration

The authors declare no conflict of interest.

## References

1. Chendrimada, T.P., Gregory, R.I., Kumaraswamy, E., Norman, J., Cooch, N., Nishikura, K. and Shiekhattar, R. (2005) TRBP recruits the Dicer complex to Ago2 for microRNA processing and gene silencing. Nature, 436, 740–744.

2. Haase, A.D., Jaskiewicz, L., Zhang, H., Laine, S., Sack, R., Gatignol, A. and Filipowicz, W. (2005) TRBP, a regulator of cellular PKR and HIV-1 virus expression, interacts with Dicer and functions in RNA silencing. EMBO Rep, 6, 961–967.

3. MacRae, I.J., Ma, E., Zhou, M., Robinson, C.V. and Doudna, J.A. (2008) In vitro reconstitution of the human RISC-loading complex. Proc Natl Acad Sci U S A, 105, 512– 517.

4. Jee, D. and Lai, E.C. (2014) Alteration of miRNA activity via context-specific modifications of Argonaute proteins. Trends Cell Biol, 24, 546–553.

5. Horman, S.R., Janas, M.M., Litterst, C., Wang, B., MacRae, I.J., Sever, M.J., Morrissey, D.V., Graves, P., Luo, B., Umesalma, S. et al. (2013) Akt-mediated phosphorylation of argonaute 2 downregulates cleavage and upregulates translational repression of MicroRNA targets. Mol Cell, 50, 356–367.

6. Zeng, Y., Sankala, H., Zhang, X. and Graves, P.R. (2008) Phosphorylation of Argonaute 2 at serine-387 facilitates its localization to processing bodies. Biochem J, 413, 429–436.

7. Rudel, S., Wang, Y., Lenobel, R., Korner, R., Hsiao, H.H., Urlaub, H., Patel, D. and Meister, G. (2011) Phosphorylation of human Argonaute proteins affects small RNA binding. Nucleic Acids Res, 39, 2330–2343.

8. Shen, J., Xia, W., Khotskaya, Y.B., Huo, L., Nakanishi, K., Lim, S.O., Du, Y., Wang, Y., Chang, W.C., Chen, C.H. et al. (2013) EGFR modulates microRNA maturation in response to hypoxia through phosphorylation of AGO2. Nature, 497, 383–387.

9. Golden, R.J., Chen, B., Li, T., Braun, J., Manjunath, H., Chen, X., Wu, J., Schmid, V., Chang, T.C., Kopp, F. et al. (2017) An Argonaute phosphorylation cycle promotes microRNA-mediated silencing. Nature, 542, 197–202.

10. Quevillon Huberdeau, M., Zeitler, D.M., Hauptmann, J., Bruckmann, A., Fressigne, L., Danner, J., Piquet, S., Strieder, N., Engelmann, J.C., Jannot, G. et al. (2017) Phosphorylation of Argonaute proteins affects mRNA binding and is essential for microRNA-guided gene silencing in vivo. EMBO J, 36, 2088–2106.

11. Agarwal, V., Bell, G.W., Nam, J.W. and Bartel, D.P. (2015) Predicting effective microRNA target sites in mammalian mRNAs. Elife, 4.

12. Krek, A., Grun, D., Poy, M.N., Wolf, R., Rosenberg, L., Epstein, E.J., MacMenamin, P., da Piedade, I., Gunsalus, K.C., Stoffel, M. et al. (2005) Combinatorial microRNA target predictions. Nat Genet, 37, 495–500.

13. Rajewsky, N. (2006) microRNA target predictions in animals. Nat Genet, 38 Suppl, S8–13.

14. Bueno, M.J. and Malumbres, M. (2011) MicroRNAs and the cell cycle. Biochim Biophys Acta, 1812, 592–601.

15. Chen, J., Cai, T., Zheng, C., Lin, X., Wang, G., Liao, S., Wang, X., Gan, H., Zhang, D., Hu, X. et al. (2017) MicroRNA-202 maintains spermatogonial stem cells by inhibiting cell cycle regulators and RNA binding proteins. Nucleic Acids Res, 45, 4142–4157.

16. Hayes, J., Peruzzi, P.P. and Lawler, S. (2014) MicroRNAs in cancer: biomarkers, functions and therapy. Trends in Molecular Medicine, 20, 460–469.

17. Peng, Y. and Croce, C.M. (2016) The role of MicroRNAs in human cancer. Signal Transduct Target Ther, 1, 15004.

18. Pan, H., Qin, K., Guo, Z., Ma, Y., April, C., Gao, X., Andrews, T.G., Bokov, A., Zhang, J., Chen, Y. et al. (2014) Negative elongation factor controls energy homeostasis in cardiomyocytes. Cell Rep, 7, 79–85.

19. Zhou, X., Geng, L., Wang, D., Yi, H., Talmon, G. and Wang, J. (2017) R-Spondin1/LGR5 Activates TGFbeta Signaling and Suppresses Colon Cancer Metastasis. Cancer Res, 77, 6589–6602.

20. Alon, U. (2007) Network motifs: theory and experimental approaches. Nat Rev Genet, 8, 450–461.

21. Cheng, C., Yan, K.K., Hwang, W., Qian, J., Bhardwaj, N., Rozowsky, J., Lu, Z.J., Niu, W., Alves, P., Kato, M. et al. (2011) Construction and analysis of an integrated regulatory network derived from high-throughput sequencing data. PLoS Comput Biol, 7, e1002190.

22. Shalgi, R., Lieber, D., Oren, M. and Pilpel, Y. (2007) Global and local architecture of the mammalian microRNA-transcription factor regulatory network. PLoS Comput Biol, 3, e131.

23. Tsang, J., Zhu, J. and van Oudenaarden, A. (2007) MicroRNA-mediated feedback and feedforward loops are recurrent network motifs in mammals. Mol Cell, 26, 753–767.

24. Martello, G., Rosato, A., Ferrari, F., Manfrin, A., Cordenonsi, M., Dupont, S., Enzo, E., Guzzardo, V., Rondina, M., Spruce, T. et al. (2010) A MicroRNA targeting dicer for metastasis control. Cell, 141, 1195–1207.

25. Tokumaru, S., Suzuki, M., Yamada, H., Nagino, M. and Takahashi, T. (2008) et-7 regulates Dicer expression and constitutes a negative feedback loop. Carcinogenesis, 29, 2073–2077.

26. Inukai, S., Pincus, Z., de Lencastre, A. and Slack, F.J. (2018) A microRNA feedback loop regulates global microRNA abundance during aging. RNA, 24, 159–172.

27. Zhang, F. and Wang, D. (2017) The Pattern of microRNA Binding Site Distribution. Genes (Basel), 8.

28. Zare, H., Khodursky, A. and Sartorelli, V. (2014) An evolutionarily biased distribution of miRNA sites toward regulatory genes with high promoter-driven intrinsic transcriptional noise. BMC Evol Biol, 14, 74.

29. Barabasi, A.L. and Albert, R. (1999) Emergence of scaling in random networks. Science, 286, 509–512.

30. Jiang, W., Guo, Z., Lages, N., Zheng, W.J., Feliers, D., Zhang, F. and Wang, D. (2018) A Multi-Parameter Analysis of Cellular Coordination of Major Transcriptome Regulation Mechanisms. Sci Rep, 8, 5742.

31. Padawer, T., Leighty, R.E. and Wang, D. (2012) Duplicate gene enrichment and expression pattern diversification in multicellularity. Nucleic Acids Res, 40, 7597–7605.

32. Davis, B.N. and Hata, A. (2009) Regulation of MicroRNA Biogenesis: A miRiad of mechanisms. Cell Commun Signal, 7, 18.

33. Shen, J. and Hung, M.C. (2015) Signaling-mediated regulation of MicroRNA processing. Cancer Res, 75, 783–791.

34. Slezak-Prochazka, I., Durmus, S., Kroesen, B.J. and van den Berg, A. (2010) MicroRNAs, macrocontrol: regulation of miRNA processing. RNA, 16, 1087–1095.

35. Darnell, R.B. (2010) HITS-CLIP: panoramic views of protein-RNA regulation in living cells. Wiley Interdiscip Rev RNA, 1, 266–286.

36. Hafner, M., Landthaler, M., Burger, L., Khorshid, M., Hausser, J., Berninger, P., Rothballer, A., Ascano, M., Jr., Jungkamp, A.C., Munschauer, M. et al. (2010) Transcriptome-wide identification of RNA-binding protein and microRNA target sites by PAR-CLIP. Cell, 141, 129–141.

37. Ke, S., Alemu, E.A., Mertens, C., Gantman, E.C., Fak, J.J., Mele, A., Haripal, B., Zucker-Scharff, I., Moore, M.J., Park, C.Y. et al. (2015) A majority of m6A residues are in the last exons, allowing the potential for 3’ UTR regulation. Genes Dev, 29, 2037–2053.

38. Chou, C.H., Chang, N.W., Shrestha, S., Hsu, S.D., Lin, Y.L., Lee, W.H., Yang, C.D., Hong, H.C., Wei, T.Y., Tu, S.J. et al. (2016) miRTarBase 2016: updates to the experimentally validated miRNA–target interactions database. Nucleic Acids Res, 44, D239–247.

39. Chou, C.H., Shrestha, S., Yang, C.D., Chang, N.W., Lin, Y.L., Liao, K.W., Huang, W.C., Sun, T.H., Tu, S.J., Lee, W.H. et al. (2018) miRTarBase update 2018: a resource for experimentally validated microRNA–target interactions. Nucleic Acids Res, 46, D296– D302.

40. Jayaram, N., Usvyat, D. and AC, R.M. (2016) Evaluating tools for transcription factor binding site prediction. BMC Bioinformatics.

41. Dermitzakis, E.T. and Clark, A.G. (2002) Evolution of transcription factor binding sites in Mammalian gene regulatory regions: conservation and turnover. Mol Biol Evol, 19, 1114–1121.

42. Berman, B.P., Nibu, Y., Pfeiffer, B.D., Tomancak, P., Celniker, S.E., Levine, M., Rubin, G.M. and Eisen, M.B. (2002) Exploiting transcription factor binding site clustering to identify cis-regulatory modules involved in pattern formation in the Drosophila genome. Proc Natl Acad Sci U S A, 99, 757–762.

